# Nested Male Reproductive Strategies in a Tolerant Multilevel Primate Society

**DOI:** 10.1101/2025.10.27.684814

**Authors:** Federica Dal Pesco, Christof Neumann, Franziska Trede, Dietmar Zinner, Julia Fischer

**Author notes:** Corresponding author: Federica Dal Pesco. **Author Contributions:** FDP, JF, and CN designed the study; FDP curated and extracted the data; CN and FDP analysed the data including the data visualisation; FT conducted the genetic laboratory analysis; DZ supervised the genetic laboratory analysis; FT and FDP conducted all analyses regarding the genotypes’ descriptive statistic and the paternity analysis; FDP, JF, and CN wrote the manuscript; all authors discussed and commented the manuscript. **Ethical approval:** All applicable international, national, and/or institutional guidelines for the care and use of animals were followed. All procedures performed in studies involving animals were following the ethical standards of the institution or practice at which the studies were conducted. Approval and research permission were granted by the DPN and the MEPN de la République du Sénégal (research permit numbers: 0383/24/03/2009; 0373/10/3/2012, 08/01/2017, 22/04/2019). Research was conducted within the regulations set by Senegalese agencies as well as by the Animal Care Committee at the German Primate Center (DPZ).

## Abstract

Male reproductive success varies within populations. Models explaining reproductive skew emphasise dominance, yet the determinants of male reproductive success in egalitarian societies remain poorly understood. Using nine years of behavioural and genetic data from five parties, we investigated male reproductive monopolisation in wild Guinea baboons (*Papio papio*), an egalitarian multilevel society with one-male “units” nested within “parties” and low male-male contest competition. Genetic analyses showed 93% of the 69 infants were sired by the female’s “unit male”, with extra-unit paternities consistent with limited control models. Reproduction was distributed among multiple unit males, resulting in low reproductive skew at the party level, with top males siring 23-40% of offspring, well below levels in despotic species. Female takeovers were rare, suggesting male restraint. Among prime-aged males, the number of unit females was only weakly positively associated with dominance ratings. Prime-age predicted reproductive success better than dominance. Non-prime-aged males had fewer females across dominance ratings. In conclusion, male Guinea baboons’ reproductive potential is shaped less by dominance than by age and stable associations with females, who play an active role in inter-sexual relationships. These results emphasise the need to move beyond frameworks focused solely on dominance-based mechanisms and to consider species-specific social organisation.

**Significance statement:** Current theoretical models of male reproductive success often focus on dominance and males’ ability to monopolise access to females. Yet, less is known about the determinants of reproductive success in societies with egalitarian male relationships, such as the multilevel society of wild Guinea baboons. Using nine years of behavioural and genetic data from five groups (“parties”), we show that unit males almost always sire their unit’s offspring. However, at the party level, male reproductive skew is low. Reproductive success is better predicted by prime-age than dominance and appears to result from long-term associations with females, likely shaped by female choice. Our results highlight the need to examine diverse social systems to understand the evolution of reproductive strategies beyond dominance-centred models.

## Introduction

Because reproductive opportunities are limited, individuals often compete for access to mates, leading to considerable variation in reproductive success. Several theoretical models have been proposed to explain patterns of reproductive skew, the unequal distribution of reproductive success. Limited control models (or ‘Tug-of-war’) suggest that reproductive skew results from reproductive contests between dominant and subordinate individuals, which impose costs to group productivity [1]. These models acknowledge that dominant individuals cannot fully monopolise reproductive opportunities, primarily due to the high costs of competition [1, 2]. Consistent with this view, the ‘priority of access model’, developed for primates, posits that higher-ranking males have priority in mating [3]. Since only one female can be monopolised at a time, the distribution of reproductive success is affected by the number of males and receptive females at a given point in time [4, 5]. Several studies have demonstrated that oestrus synchrony limits the success of male monopolisation in primates, highlighting how breeding seasonality and the number of females in a group influence male reproductive skew [5–7]. Transactional models, in contrast, assume that reproduction can be controlled and suggest that males share reproduction through concessions or restraint to maximise group stability and mutual benefits [8–10]. The magnitude of concession or restraint is determined by kinship and other social and ecological factors [11].

Variations in the degree of male-male contest competition and dispersal patterns influence the outcomes of conflicts over reproduction. In societies where direct male competition is reduced and hierarchies are less strictly enforced, males may engage in reproductive cooperation through alliances and bonds that provide both indirect and direct fitness benefits [e.g., 12, 13]. Male-philopatric societies, characterised by higher relatedness among males, are expected to have low to moderate levels of reproductive skew, most likely due to kin-based alliances and cooperation among males [discussed in 14]. A striking example is the patrilocal society of northern muriquis (*Brachyteles hypoxanthus*) where aggressive competition between males is virtually absent and male reproductive skew extremely low [14]. Bonobos (*Pan paniscus*) also live in male philopatric societies, yet high-ranking males successfully monopolise a large share of reproduction, with the most reproductively successful male siring up to 62% of offspring [15]. Interestingly, reproductive skew in bonobos exceeds that of chimpanzees (*Pan troglodytes*), a closely related species that also lives in fission-fusion patrilocal societies but presents more frequent and intense aggression between males [15]. In chimpanzees, reproductive skew was lower but varied substantially between communities, likely reflecting environmental and demographic factors [12]. The stronger association between dominance rank and reproductive success in male bonobos, despite less intense mate competition, has been attributed to greater group cohesiveness, which may facilitate monopolisation, and to female choice for high-ranking males in these female-dominated societies [15].

In many mammals male age also plays a key role in shaping reproductive monopolisation, as it is linked to physical maturation, body size, and the development of sexually selected traits [16]. In most species, subadults have lower reproductive success compared to adults [17, 18] and prime-age males, being at their peak physical condition and highest rank [16], have higher reproductive success compared to other age classes [18]. In some species, however, older males can be equally or even more successful. In African elephants (*Loxodonta africana*), for example, male rank depends on body size, and males grow throughout much of their lives [19], with older males having markedly elevated paternity success [16].

Male reproductive success depends not only on the ways males attempt to gain access to mates, but also on female behaviour. In spotted hyenas (*Crocuta crocuta*), where females typically dominate males, dominance and direct competition among males have little effect on male reproductive success, which is primarily affected by female mate choice [20]. In vervet monkeys (*Chlorocebus pygerythrus*), where females exhibit co-dominance, females mate with multiple males and can resist mating attempts, influencing reproductive monopolisation [21]. In several species, including some in which males are dominant over females, female preference for males that provide support and protection is associated with variation in male reproductive success [22–26].

To date, the determinants of male reproductive success in egalitarian societies, characterised by low levels of contest competition, shallow dominance hierarchies, and high social tolerance, remain less well understood. To fill this knowledge gap, we investigated male reproductive monopolisation in wild Guinea baboons (*Papio papio*), a species living in a multilevel society with female-biased dispersal [27, 28], egalitarian relationships among males [29, 30], and relatively high female leverage [31, 32]. The core of the society consists of ‘units’ comprising one reproductively active ‘primary’ male (hereafter ‘unit male’), typically 1-8 associated females, and their offspring ([31, 32]; unpublished data). Male-female associations remain stable throughout female reproductive cycle and can last from a few weeks to several years ([31], unpublished data). Several units and bachelor males associate in stable social groups called ‘parties’ (intermediate level), where males maintain strong bonds and display a high degree of social tolerance [27, 30, 32]. Bachelor males may associate with one or, in most cases, several unit males and play an integral role in party cohesion [30]. Several parties form the upper-level social grouping known as a ‘gang’, and several gangs constitute the “community” (apex level). These gangs share largely overlapping home ranges [27, 33], highlighting the high levels of spatial tolerance of Guinea baboons both within and between social levels [reviewed in 34]. This multilevel organisation is similar to that of other papionins, such as geladas (*Theropithecus gelada*) and hamadryas baboons (*Papio hamadryas*), which are characterised by stable core units embedded within higher-order aggregations [35]. However, beyond this shared architectural template, these species differ in dispersal patterns and the nature of social relationships within and between levels [35, 36].

In Guinea baboons, the party constitutes the stable and visibly cohesive intermediate level of the society and is the primary context in which male–male social relationships are maintained [27, 30, 37]. The party is the relevant level at which males can be compared. Units, by definition, comprise only one reproductively active male, rendering comparisons between males moot. Within parties, males form strong, lasting bonds with other males and rely on them for mutual support during aggressive events [32]. Strong ties between males, however, are not linked to increased reproductive success. Instead, as the number of females in their units increases, males tend to spend less time socialising with other males [32].

The rate of aggression between Guinea baboon males is lower than in other Papio species [29] and their dominance hierarchies were previously described as “non-significant” [29] and characterised by intermediate/low steepness [30]. In line with this generally low level of overt male–male contest competition, and given the scarce occurrence of decided agonistic interactions between males, previous analyses have failed to estimate male dominance hierarchies with any certainty [30], raising the possibility that rank is of little utility in describing male-male relationships. Similar patterns have been reported in other multilevel societies, where male status (e.g., rank) is often more closely linked to social roles (e.g., leader/follower) or age classes than to clearly expressed linear dominance hierarchies [e.g., 38, 39]. Alternatively, existing methods [40] may not have been able to estimate reliable dominance hierarchies under such conditions. To address these two possibilities directly, we employed novel statistical methods that explicitly incorporate uncertainty during the estimation of individual dominance ratings (‘ranks’) and utilise such ratings, along with their associated uncertainty, as predictors in our modelling approach. Estimating dominance ranks, ratings, and hierarchies is challenging, especially when few decided interactions were recorded (due to low interaction rates and/or low observation effort) and/or under conditions of low hierarchy steepness [40]. With this novel approach, we aimed for a more nuanced understanding of social and reproductive dynamics in this species. Rather than assuming that rank is or is not biologically meaningful a priori, or inferring its relevance from the rate of agonistic interactions alone, we aimed to empirically test whether male rank was associated with variation in reproductive opportunities, while accounting for uncertainty in rank estimation.

We used long-term data on wild Guinea baboons collected over nine years from five social groups (parties) to investigate the extent of male reproductive skew and the link between male rank and age and their reproductive success. The intermediate level of the party was used as our group unit because males aggregate and interact the most at this social level [27, 30]. We first established paternity for all genotyped offspring. Considering the high investment of unit males in stable year-round associations with their females and the high mating fidelity [99% of observed copulations, 31], we predicted high paternity certainty (i.e., unit males should sire the offspring of their associated females). We then assessed reproductive skew by quantifying the distribution of reproductive benefits among all co-resident males at the party level. Given the multi-level social organisation of the species, the presence of male kin in the party, and the occurrence of male-male social tolerance and bonds, we hypothesised that the balance between competition and cooperation is shifted towards male cooperation, resulting in low reproductive skew. Lastly, we modelled the effects of age and rank on male reproductive success using a custom Bayesian model based on Elo-ratings, while propagating uncertainty in the rank estimates into our model of reproductive opportunities. Given that Guinea baboons exhibit low rates of overt male-male competition, we predicted that male rank would have a modest linear effect on reproductive success. We further predicted that males in their prime age have a higher reproductive success than subadult males or males past their prime.

## Material and methods

### Study subject and study site

We used data collected over nine years (2014-2022) on wild Guinea baboons at the CRP Simenti field station in Niokolo-Koba National Park, Senegal [34]. We initially worked with three habituated parties (5, 6, 9) within the Simenti Guinea baboon community, which consists of around 400 individuals. Party 6 split into two parties, 6I and 6W, in 2020. Ultimately, we entered five party IDs into our analyses (see supplementary table S1 for details). We confined all our analyses to within-party interactions, following previous studies on the same population [e.g., 30, 32]. We conducted behavioural observations on all males in the study parties, starting from the time they first attained the age classification of large subadult, approximately 8 years old (*n* = 48). At that age, they may already be associated with females and hold their unit [see 32]. For each study year, we assigned males to one of two age categories. Sexually mature and fully developed individuals were categorised as prime-age. Individuals not yet fully developed, as well as old males, were classified as non-prime-age. For males who switched from non-prime to prime-age or vice versa in the same study year, the category with the longer duration (higher number of days) was chosen for that year. We collected data using customised electronic forms, which were operated on Pendragon 7.2 software, and were accessed via Samsung Note 2 and 3, as well as Gigaset GX290 Plus handheld devices. On each observation day, we recorded census data encompassing demographic information (including births, deaths, dispersal/migration, and presence/absence), health status, and female reproductive state [see 31]. We collected 20-minute continuous focal follows [41], ensuring a balance between subjects and the time of day. During these focal follows, we continuously recorded the focal animal’s activity (movement, feeding, resting, and socialising) and all social behaviours (e.g., approaches, retreats, supplants, aggression, grooming, contact-sits, and greetings) involving the focal subject. Instances of aggression, coalitionary support, copulation, and grooming were also recorded on an ad libitum basis.

### Number of associated females (unit size)

We recorded unit composition and female unit transfers on each observation day. Daily unit composition was established based on previous observations showing that females interact more with their unit males [31]. We thus based our assessment of unit composition on copulations, grooming bouts, contact-sit bouts, greetings, and aggression events between each female and all males in the same party [see 32]. To account for unit size variation due to female transfers and demographic changes, we used daily unit size within each study year to calculate the modal number of associated females per male and year (i.e., the most frequent unit size value), following [32]. The daily unit size for bachelor males was, by definition, zero.

### Genetic and paternity analyses

We used genetic data collected over eight years (2014-2021) to evaluate male paternity, reproductive skew, and the proportion of offspring sired by males of the five study parties. A subset of these data (2014-2017, two parties) was previously analysed by [32] to investigate the fitness benefits of male-male sociality. The present study expands this dataset by including additional genetic and demographic data collected from 2018–2021 and from additional parties (see supplementary table S2). All samples were collected and analysed using the same standardised field and laboratory protocols described below and in [32].

During 2014-2021, 110 infants were born within the study parties, of which 71 were sampled successfully for genetic analysis (supplementary table S4). To conduct the parental analysis, we utilised faecal samples from all focal males (from approximately 8 years old) and sexually mature females (from approximately 5.5 years old) who were members of the study parties during the period of interest. We accounted for a 6-month gestation period (see supplementary table S2). This value is based on long-term demographic data from the study population, including observed conception–birth intervals in the dataset analysed here, which indicate a median gestation length of approximately 5.8 months (unpublished data). We used a rounded value as a conservative approximation given within-population variation in gestation length. Since we did not know at which age sperm would be viable in our population, and given that in other baboon species, even relatively young males have been documented to sire offspring [42], we also included all males between 6 and 8 years of age. Ultimately, we sampled 54 males (98% sampling success, with all unit males present at the time of conception sampled) and 68 females (85% female sampling success, with all observed mothers sampled). In addition, to rule out potential cases of paternity outside of the study parties or errors in our maternity assessment, we collected faecal samples from 50 males and 13 females belonging to the other habituated parties within our study population.

We assessed individual allelic variation using a panel of 24 autosomal microsatellite markers [43, also see 27]. The methods employed in genetic sample collection, storage, DNA extraction, and genotyping are described in Dal Pesco et al. [43]. All loci demonstrated polymorphism, with an average allele number of 4.5 (range = 2-7; sd = 1.4); no locus showed signs of null alleles or departures from Hardy-Weinberg Equilibrium (see supplementary table S3).

The paternity analysis was performed using the software Cervus [v. 3.0.7, 44], employing the methodologies described in detail by Dal Pesco et al. [43]. The mother’s identity was recorded during field observations and verified using a maternity likelihood analysis with the acceptance criterion of identification as candidates with zero mismatches. All mothers were confirmed with zero mismatches. Subsequently, we employed the trio likelihood approach, leveraging the known identity of the mother to determine the most probable father (supplementary table S4). A male was considered the sire of an offspring if he was assigned as the most likely father, had a maximum of one mismatched allele, and the confidence level for the assignment exceeded 95% (using the ‘strict’ criterion).

### Reproductive skew and male siring success

For each genotyped offspring born between 2014 and 2021 (*n* = 71), we checked the unit association of the mother at the time of conception. Then we calculated the percentage of offspring sired by each unit male. For two offspring, we did not know with which male the female was associated before they joined our study groups, resulting in 69 offspring for which we could determine whether the unit male sired the offspring (see supplementary table S4).

To measure male monopolisation at the party level, we calculated the percentage of offspring sired by the most successful male within each party over the eight years. Initially, we calculated this percentage using only the offspring sired within the party for which paternity was genetically assessed (*n* = 70, one case of extra-party paternity was excluded). Based on high paternity certainty (see results), we assigned the unit male at the time of conception as the most likely father for offspring without genetic data. We then repeated the calculation, including all offspring sired within the party (*n* = 109).

To investigate male reproductive skew at the party level, we used all offspring with assigned paternity in the study parties (*n* = 70). One offspring was sired by a male of another party (see supplementary table S4). We included all focal males associated with the study parties over the eight years, extending from six months before the start to six months before the end of each study party’s observation period to account for gestation time (supplementary table S2). This dataset comprised 45 males (note that one male was counted twice due to a party transfer). Although Party 6 split into Parties 6I and 6W in 2020, we combined them for the reproductive skew analysis and for calculating the share of offspring sired by the most successful male, because the split occurred near the end of genetic data collection and sample sizes were too small for separate analyses. All other analyses treated these parties separately after the split.

We calculated reproductive skew using the Multinomial Skew Index [*M* index, 45, see supplement section “Reproductive skew” for details about implementation and interpretation]. This index can be fitted in a Bayesian framework and therefore results in a posterior distribution of M. In contrast to the widespread B index (which can still be calculated from M), M is unconstrained. An M value equal to zero indicates that reproduction is distributed as expected for a random multinomial model where rates of reproductive success are equal. An M value above zero indicates a positive skew of reproduction. An M value below zero indicates that the share of reproduction within the group is more equal than expected under a random multinomial model [45]. The share of reproduction is not different from chance when the confidence interval includes zero. The M index accounts for group demographic changes, allowing us to include demographic information regarding male presence, and is not biased by differences in sample or group size [45]. We also calculated M for the total monopoly and equal sharing scenarios [46] to assess where these values fall in relation to the observed indices (see supplement section “Reproductive skew”).

### Dominance hierarchy

We evaluated the male dominance hierarchy using ad libitum and focal data on within-party male-male aggressive behaviours that occurred during the nine-year study period. This included all threats, lunges, chases, physical fights, displacements/supplants, avoidance, and unprovoked submissions. We excluded interactions without a decided winner/loser outcome. We also excluded all interactions that followed one or more polyadic interactions within the same aggression event, as the outcome could still be influenced by previous coalitionary behaviour. In cases where multiple interactions occurred between dyads during the same aggression event, we only considered the most intense interaction. We included 714 interactions in the rank calculation (median = 33.5 per male, range = 1 *−* 90). We assessed male rank within parties by estimating Bayesian Elo-ratings [47, see supplementary subsection “Elo-ratings” for a full methodological description].

### Modelling potential reproductive success as a function of age and rank

We modelled the modal number of females a male had in his unit in a given calendar year and use this response as a proxy for potential reproductive output [see 32]. Our data set comprised 190 data points (male-years), with 48 male subjects. Males who switched party association within a calendar year were assigned to the party in which they resided the longest. Males were present between 1 and 9 years (median = 3). We modelled the modal number of females a male had in his unit in a given year as a Poisson process with a log-link. As predictors, we used a male’s Elo-rating (at the end of a given calendar year), age, and the interaction between Elo-rating and age. We fit the model in a Bayesian framework in Stan [48, 49], which allowed us to combine Bayesian estimation of Elo-ratings with our model of interest on the number of females. This approach allowed us to utilise the full posteriors of Elo-ratings as a predictor variable. A detailed model description can be found in the supplementary section “Modelling potential reproductive success”.

## Results

### Reproductive skew and male siring success

We were able to estimate paternity for all 71 sampled offspring. Of these, 67 offspring were sired by adult males, while four different subadult males sired the remaining four offspring. Of the 69 offspring for which we could determine the identity of the unit male at the time of conception, 64 (93%) were sired by that male. The remaining five offspring were sired by an adult male from a different party (same gang), whose social association with the mother ended 9.5 months prior to birth (i.e., before the likely conception window), and by four subadult males belonging to the same party that had bachelor status at the time of conception (see supplementary table S4). The units in which these five extra-unit paternities occurred contained two or three females at the time of conception. All extra-unit paternities occurred between August and November, corresponding to the middle or late rainy season, or shortly thereafter, when the vegetation is dense and visibility is poor (not only for the observers but most likely also for the baboons). For all extra-unit paternity cases, we examined whether any other unit females were cycling simultaneously during the conception period. For the four cases for which we had these data, the unit male had two females cycling at the same time during the conception period.

At the party level, reproductive success was distributed across multiple males, predominantly the female’s unit male at the time of conception. If we exclude the one extra-party paternity, the remaining 70 offspring had 20 different fathers, indicating that at the party level reproduction is distributed among multiple males. Almost half of the large subadult and adult males of the study parties (47%) sired at least one offspring. The number of sired offspring per male during the study period ranged from 0 to 8 (mean = 1.56; sd = 2.23). When we considered all offspring sired within the party, regardless of whether they were genotyped, the 109 offspring had 27 different fathers, with 60% of males siring at least one offspring (mean = 2.42, sd = 3.09, range = 0–11).

The percentage of offspring sired by the most successful male within each party ranged from 23% to 40% over the eight-year study period (average across parties 29%) when considering only offspring for which paternity was genetically assessed; in two of the three parties, two or three males tied as the most successful. When including all offspring, this percentage ranged from 24% to 33% (average across parties 28%), and each party then had a single most successful male. The expected distribution under random paternity (i.e., assuming all males have equal probability of siring each offspring), given the number of candidate males per party, would range between 6% and 11% (average across parties 7%), indicating that the observed distributions are modestly higher than expected by chance. These values reflect the distribution of realised paternities among males within parties and are therefore calculated at the party level, rather than being adjusted for variation in unit composition or number of females associated with individual males. This approach allows us to assess how reproduction is distributed among the multiple males co-residing within a party, beyond the high concentration of paternity within individual units.

The reproductive skew, measured within parties using the multinomial index, was close to zero, with posterior medians ranging from 0.24 to 0.54 for the three parties (table 1). The 89% credible intervals overlapped zero for all parties, indicating that the distribution of reproductive benefits within parties (i.e., sired offspring in the study parties) did not differ from chance in an obvious way (figure 1). For all parties, the point estimate for the total monopoly scenario was well outside the credible intervals (table 1), providing strong evidence that reproduction is not monopolised by a single male at the party level but rather that multiple males, most often unit males, reproduce. The point estimate for the equal share scenario was just outside the confidence interval ranges for all parties (table 1), indicating that reproduction is not entirely distributed equally. Inequalities likely emerge from differences in male status and unit size.

**Table 1.** Reproductive skew assessed by the multinomial index (M) in three parties of Guinea baboons. We investigated reproductive skew for all focal males and all offspring present/born between 2014 and 2021 with assigned paternity in our three study parties.

| party | males | sires | offspring | $M_{\text{post. median}}$ | 89% CI | monop | equal |
| --- | --- | --- | --- | --- | --- | --- | --- |
| 5 | 9 | 5 | 20 | 0.33 | -0.39 – 1.61 | 7.60 | -0.90 |
| 6 | 18 | 10 | 28 | 0.54 | -0.43 – 2.05 | 16.39 | -0.94 |
| 9 | 18 | 6 | 22 | 0.24 | -0.85 – 2.02 | 16.22 | -0.95 |

**Figure 1.**
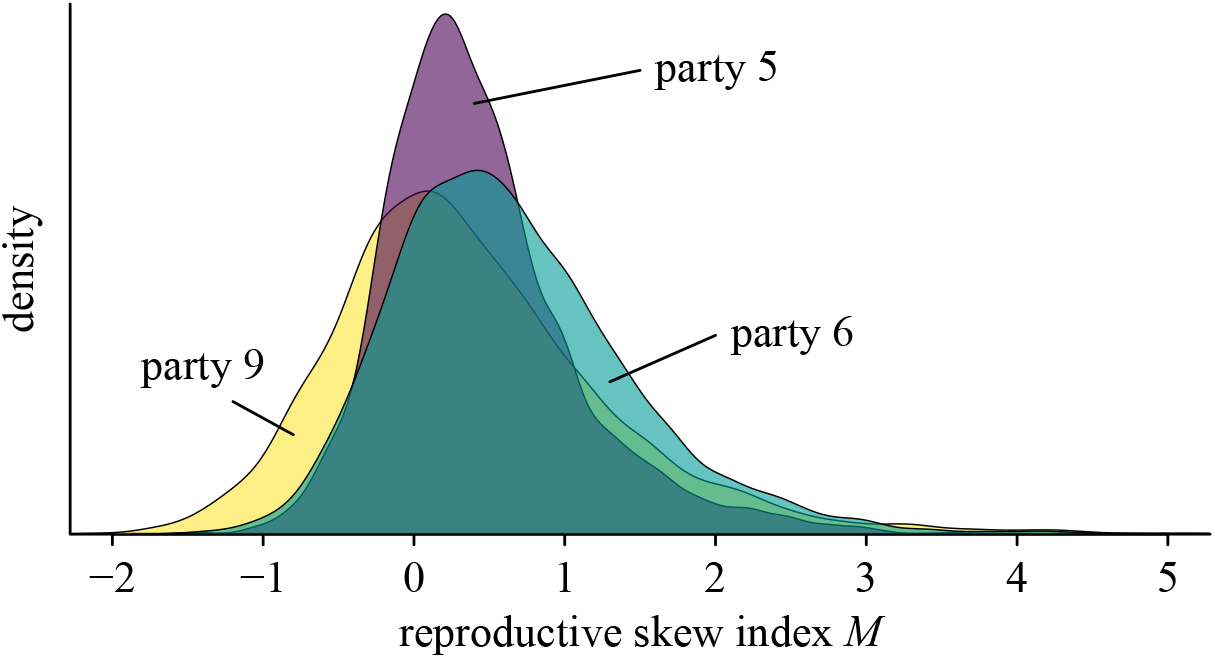
Full posterior distributions of the reproductive skew index for three Guinea baboon parties (5, 6, and 9).

### Modelling potential reproductive success as a function of age and rank

We found a positive relationship between a male’s Elo-rating in a given year (figure 2) and the mode number of females a male was associated with in that year (a proxy for potential reproductive output, given high within-unit paternity certainty) (figure 3, table 2, see supplementary figure S6 for result visualisation by party). The positive relationship between rank and reproductive success was only visible in prime-aged males. In contrast, for non-prime-aged males, the association between rating and number of females was virtually flat (see figure 2 for a visualisation of Elo-rating ranges by age). Reproductive success was not biased in favour of the highest-ranking males. Indeed, middle-ranking males held the largest units (4-6 females), and some of the top-ranking males had no units or units with only 1 to 2 females. Across males and years, the average mode of the number of associated females was 1.78 (*n* = 148 male-years, sd = 1.42, range = 0-6) for prime-aged males and 0.14 (*n* = 42 male-years, sd = 0.42, range = 0-2) for non-prime-aged males. Of the 148 male-year observations for prime-aged males, only 24% had a mode number of females of zero (*n* = 20 males). Non-prime-aged males typically had no females and there were only five observations where non-prime-aged males had any females; these were large subadult males.

**Figure 2.**
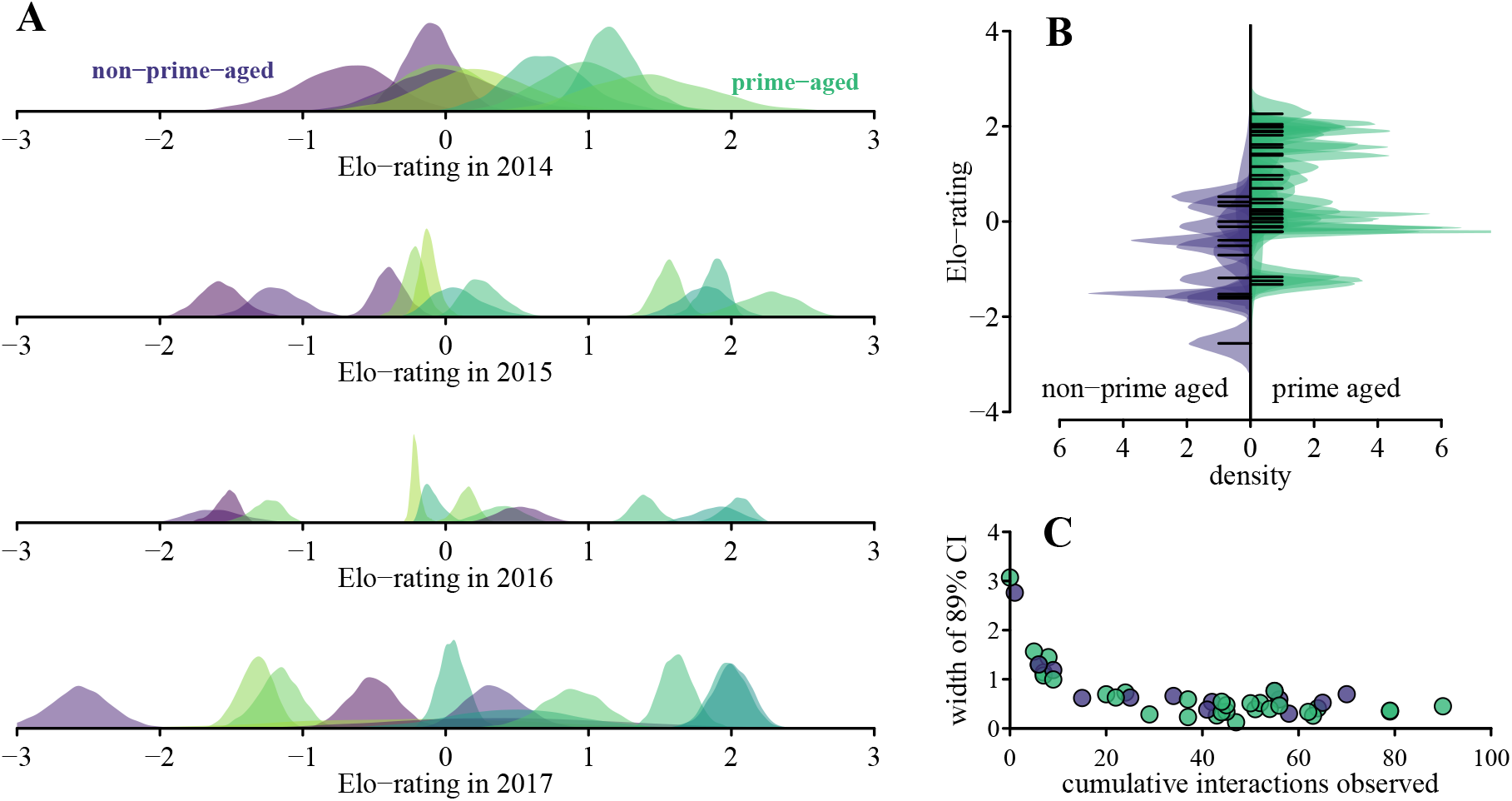
Bayesian Elo-rating. We assessed male rank within parties using a modified Bayesian Elo-rating approach that yields ratings as posterior distributions instead of conventional point estimates. In our down-stream model of reproductive success, we make use of these full posteriors as predictors, thereby carrying forward the uncertainty from rank estimation. The three panels illustrate this approach for a sub-sample of four years and one study party (party 6). Panel A represents the individual dominance ratings and associated posteriors at the end of each calendar year. Panel B shows the range of rating posteriors across age classes. Panel C illustrates how uncertainty in the rating estimates typically decreases with increasing numbers of interactions.

**Figure 3.**
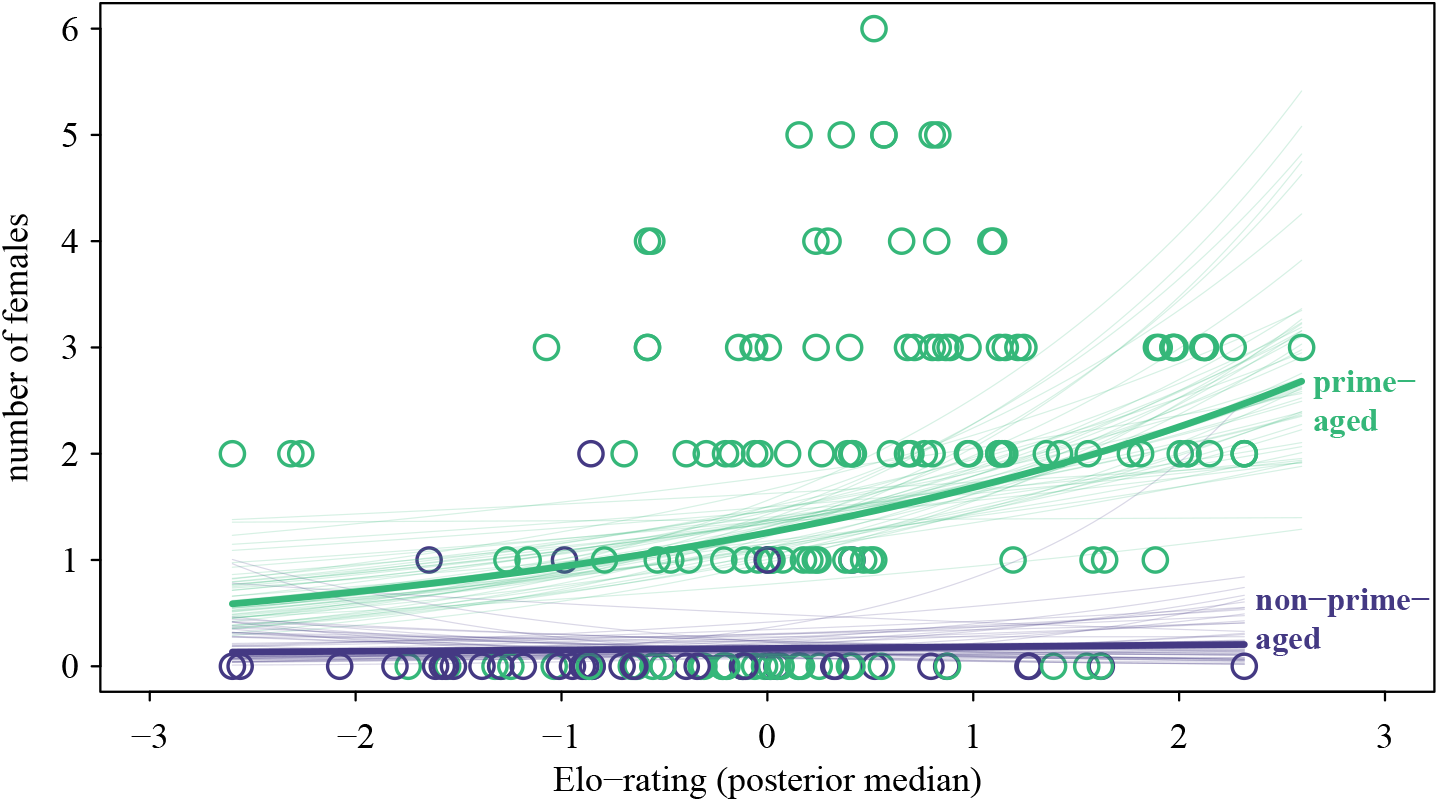
Count model results. The figure shows the mean predictions for the relationship between Elo-rating and the number of females, separated by age class. Alongside the mean predictions, 50 randomly selected samples show the range of uncertainty in the estimation. Circles represent the observed number of females, and Elo-ratings are plotted at the median of the posterior distribution for each male year.

**Table 2.** Summaries of estimates of primary model parameters, including 89% credible intervals. Greek letters refer to supplementary equation S10. The model parameters in this table represent population-level effects. Age was a binary predictor with the reference level being prime-age.

| | mean | median | 5.5% | 94.5% | $\hat{R}$ |
| --- | --- | --- | --- | --- | --- |
| intercept ($\alpha$) | 0.23 | 0.23 | -0.04 | 0.48 | 1.00 |
| Elo-rating ($\beta$) | 0.29 | 0.29 | 0.10 | 0.49 | 1.00 |
| age ($\gamma$) | -2.02 | -2.00 | -2.73 | -1.33 | 1.00 |
| interaction Elo-rating : age ($\delta$) | -0.21 | -0.21 | -0.78 | 0.38 | 1.00 |

## Discussion

Using nine years of behavioural and genetic data from five parties, we examined how reproductive success was distributed across males at different social levels of a wild Guinea baboon society. The paternity analysis revealed that 93% of the offspring were sired by the female’s unit male, indicating high paternity certainty for these males. This finding is in line with the observation that 99% of observed copulations occurred with the unit male [31] and with findings in other primate multilevel societies where unit males monopolise the majority of mating opportunities [13, 50]. Reproductive concessions to associated bachelors in exchange for support, however, appear unlikely in Guinea baboons, given that bachelor associations span multiple units and that extra-unit paternities can occasionally occur across parties. Extra-unit paternities were rare and occurred when multiple females from the same unit cycled simultaneously and during periods of low visibility due to dense vegetation, when male visual control over female whereabouts was low. We found no indication that extra-pair paternities occurred more prominently in particularly large units, as all of the five observed cases occurred in units with two or three females. This constellation of factors likely limited the unit males’ ability to completely monopolise reproduction [5, 6, 51, 52].

While these findings demonstrate strong reproductive monopolisation within units, they do not tell us how reproductive success is distributed among the multiple males that coreside within a party. The party constitutes a stable and visibly cohesive social level of Guinea baboon society and, because each unit comprises a single reproductively active male, provides the primary context in which multiple males interact and their reproductive success can be compared [27, 30, 37]. At the party level, reproduction is distributed across multiple males. About half of the males (47%) sired at least one offspring, and the most successful males accounted for 23–40% of paternities. Thus, although paternity was strongly concentrated within units, reproduction was not strongly concentrated in a single male at the party level. The most successful males account for substantially less reproduction than previously reported in some strictly hierarchical or coercive species, such as mountain gorillas living in multi-male groups (*Gorilla beringei beringei*: 85%; [53]) or yellow baboons (*Papio cynocephalus*: 81%; [54]). At the same time, this pattern is comparable to other primate multi-level societies, in which reproduction is strongly concentrated within units but distributed among multiple unit males at higher social levels [13, 50].

Our analysis indicates that the same social system can conform to predictions from different bodies of theory, depending on the social level. At the unit level, findings are consistent with limited control models where monopolisation is limited by reproductive synchrony and male visual control [1, 9, 51]. At the party level, in contrast, the combination of low reproductive skew, rare takeovers [31], and little overt aggression [30] is consistent with models of male restraint [10], whereby males limit overt competition with one another in ways that may facilitate party cohesion. This pattern resembles what Kummer described as the “respect of ownership” in hamadryas baboons [55, 56]. Taken together, these findings show that in multi-level societies, different theoretical frameworks are not mutually exclusive, but rather complement each other.

While nested social levels are a key feature of multilevel societies, most animal groups are also internally substructured and embedded in larger collectives that face similar ecological and social problems. Many theoretical models of reproductive skew assume that a single level within social groups represents the main arena for reproductive competition. We argue that explicitly considering social substructuring can help connect theoretical models developed largely from simpler social systems with the more complex organisation of multilevel societies, where different processes may operate at different social levels. The effect of substructures on social behaviour is not restricted to Guinea baboons: Recent experimental studies and agent-based modelling approaches in humans have shown that the permeability of group boundaries and the frequency of interactions between subgroups determine whether cooperation remains group-bound or transitions into broader collective actions [57]. Cross-group interactions also shift perceptions of group membership, with individuals increasingly identifying with the collective rather than their in-group [57]. Similar considerations may also apply to bonobos and dolphins, where interactions across subgroups and the permeability of social boundaries can influence patterns of cooperation, competition, and resource access [58].

In addition to understanding evolutionary dynamics, substructuring needs to be considered in comparative research. When the social level at which reproductive competition occurs is not explicitly considered, comparisons across species risk conflating fundamentally different underlying processes. We propose that the Guinea baboon “party” constitutes the ecological equivalent to the “group” in uni-level societies [36]. Thus, in the Guinea baboon male philopatric society, units comprise a single male, while the intermediate level of the party, composed of multiple unit males, their units, and bachelor males, constitute the main social arena in which males interact and establish their social relationships [30]. Considering this social level may therefore provide a more meaningful basis for comparison with species in which males interact and compete within a single cohesive group.

In prime-aged Guinea baboon males, dominance ratings were positively associated with the number of females in the unit, although the association was weak. The majority of prime-aged males had one to three females. Males with average dominance ratings had the most females, while some males with above-average dominance ratings had few or no associated females. While this pattern may question the application of a linear model, the choice of the linear model was based on the theoretical considerations outlined in the introduction. Overall, male age was a stronger predictor of reproductive success than rank. Most non-prime-aged males had no associated females, while prime-aged males typically had more females across the entire range of rank ratings. These results suggest that we are missing an important component in the model, most likely female preferences.

Females may prefer males that are more tolerant and affiliative, socially compatible, or better fathers [31, 59], even when these males hold average dominance ranks. Familiarity might provide another plausible explanation, as females might prefer to remain with males they already know well, while the composition of females within a unit may further influence mate choice. Male physical condition and developmental trajectories, such as the transition speed to adulthood, may serve as critical cues for females. Males typically establish their unit during the final stages of subadulthood, and most prime-aged males are unit males. Males then gradually lose their associated females as they transition from late prime to old age [32]. Lifetime reproductive success thus likely depends on the total number of female-years a male accumulates. More data on complete male life-histories will be needed to test this assertion.

Maximising the number of female-years aligns with patterns observed in other primate multi-level societies. The underlying behavioural strategies, however, can differ substantially between species. In hamadryas baboons, male access to females is most often achieved via coercive tactics and enforced proximity, which are thought to have evolved due to their direct fitness benefits [60–62]. Male hamadryas baboons protect their units through both direct contest competition during takeovers and alliances with follower males, which are exclusively associated with their unit [13]. In these male-philopatric societies, followers are thought to diminish the likelihood of extra-unit paternity and benefit from indirect fitness benefits and/or the possibility of future succession [13]. Their presence is also associated with increased reproductive success, with a greater number of infants born into units that include follower males [13]. In snub-nosed monkeys (*Rhinopithecus roxellana*), cooperative defence of females occurs across units. Unrelated unit males join forces against satellite males, who, in turn, cooperate in takeover attempts [63, 64]. In geladas, the most successful unit males are those that associate with follower males, benefiting from their support [50]. These follower males, in turn, gain reproductive opportunities, making geladas one of the few examples supporting transactional concession models in primates [50, but see 65]. In conjunction, depending on ecological conditions, evolutionary history, and dispersal patterns, different processes may lead to similar-looking outcomes at the level of the social structure.

Dominance rankxs has long been used as a proxy for competitive ability and mating access in group-living mammals [66], yet our results show that substantial variation in male reproductive success cannot be explained solely by dominance-based monopolisation or overt contest competition. Even among prime-aged males, individuals with similar rank differed markedly in reproductive outcomes, such that key components of fitness variation remain unaccounted for. This extends beyond Guinea baboons. Alternative pathways to reproductive success, including the role of female spatial and social preferences, long-term social relationships, and coalitionary dynamics, have been reported for other species [e.g., 20, 21, 22, 24]. While alternative pathways may remain obscured in despotic societies, expanding the mechanisms considered in reproductive skew models reduce the risk of overlooking key sources of fitness. These implications are not only limited to egalitarian societies. Even in species with clear dominance hierarchies, such as Barbary macaques (*Macaca sylvanus*), yellow baboons, and chimpanzees, reproductive outcomes can be shaped by rank, coalitionary support, or female mating behaviour rather than rank alone [26, 67–70]. In this sense, our results do not represent an exception to prevailing theory; rather, they reveal where current theoretical models are incomplete. Incorporating these dynamics into comparative analyses may help reconcile inconsistent relationships between rank and fitness across taxa.

Our study also highlights the importance of considering uncertainty when estimating social features, such as dominance scores. By conventional standards [e.g., 40], our observed interaction networks were too sparse to reliably estimate individual dominance ratings as point estimates [see 30], raising the question of what group members can infer from such rare agonistic interactions. Here, we built on recent advances in Bayesian modelling of interaction data [e.g., 47, 71, 72], which explicitly incorporate uncertainty and allow estimates to become more precise as observation records accumulate. We extended this approach by implementing a custom Stan model that directly incorporates dominance ratings and their associated uncertainty as a predictors in our statistical model of reproductive success. This approach enabled us to retain and model the uncertainty in Elo-rating, one of the two key predictors in our model, which decreases as the number of observed interactions per individual increases (see panel C in figure 2), rather than treating the estimated ratings as fixed point estimates. The key advantage of this approach is that predictors associated with uncertainty provide a more honest reflection of the underlying variable, which is the foundation of measurement error models [73].

Our results therefore provide an empirical test of whether variation in agonistic asymmetries is associated with reproductive opportunities, rather than establishing a priori that rank scores represent either a biologically meaningful hierarchy or an irrelevant construct. We found only a weak association with unit size, suggesting that rank is unlikely to be a major determinant of the reproductive opportunities examined here. Whether these agonistic asymmetries influence other aspects of male social behaviour or fitness, or whether males themselves perceive and respond to them as a social hierarchy, remains an open question.

In summary, dominance and fighting ability play only a limited role in Guinea baboons’ mating outcomes. We hypothesize that reproductive success is primarily influenced by long-term male-female associations rather than by male dominance, coercion, or coalition-based competition. Males strike a balance between competitive ability and social investment toward females, making them more attractive partners. Unlike other multilevel primate societies, where male reproductive success is achieved through aggression or strategic alliances [13, 60, 64], Guinea baboons exhibit a system that revolves around social tolerance and female choice. Our results underscore the need to move beyond dominance-based frameworks when studying reproductive skew, highlighting the role of female choice in shaping male reproductive success and the importance of considering species-specific social organisations when evaluating reproductive strategies. We suggest that two levels of reproductive strategies exist within the Guinea baboon society: limited control within units and restraint among males at the party level. Considering the effects of substructuring on social dynamics constitutes an exciting and challenging avenue for future research.

## Supporting information

SOM

## Acknowledgements

We would like to thank the Diréction des Parcs Nationaux (DPN) and Ministère de l’Environnement et de la Protéction de la Nature (MEPN) de la République du Sénégal for permission to work in the Niokolo-Koba National Park. We particularly thank all the conservateurs of the park during the study period (Oussoumane Kane, Mallé Gueye, Amar Fall, Assane Ndoye, and Jacques Gomis) for their cooperation and logistical support. We thank all the staff, field assistants, and students of the CRP Simenti for their precious support and work in the field. In particular we are grateful to all of the park agents affiliated to the CRP Simenti project for their invaluable support during daily field work: Cheikh Sané, Moustapha Dieng, Moustapha Faye, Armel Louis Nyafouna, El’Hadji Yankhoba Dansokho, Touradou Sonko, Vieux Biaye, Djibril Coly, Amadou Bamba Diedhiou, and Chérif Younousse Kéba Camara. We are grateful to Cody T. Ross for help with the SkewCalc package and the analysis of reproductive skew. This research was supported by the Deutsche Forschungsgemeinschaft (DFG, German Research Foundation) - Project number 254142454 / GRK 2070 and Project-ID 454648639 SFB 1528 „Cognition of Interaction”.

