## Supplementary material for "Nested Male Reproductive Strategies in a Tolerant Multilevel Primate Society": SOM

### S1 Study parties and subjects

Table S1: Overview of behavioral observation periods for each study party

| Party | Start | End |
|---|---|---|
| 9 | 2014-04-01 | 2020-04-12 |
| 5* | 2016-01-01 | 2022-12-31 |
| 6 | 2014-04-01 | 2020-03-31 |
| 6I* | 2020-04-01 | 2022-12-31 |
| 6W* | 2020-04-01 | 2022-12-31 |

* Covid-19 Gap: no data from 2020-04-13 until 2020-11-29

Table S2: Overview of periods of available demographic and genetic data for each study party. The columns offspring start/end indicate the periods in which newly born offspring were assessed; the columns parents start/end indicate the periods used to establish the presence of subadult females/males in the study parties as potential mothers/fathers. Note that parental presence was assessed taking into account gestation time; thus, we considered the period starting from six months before the "offspring start" date and ending six months before the "offspring end" date.

| Party | Offspring start | Offspring end | Parents start | Parents end |
|---|---|---|---|---|
| 9 | 2014-01-01 | 2020-04-12 | 2013-07-01 | 2019-10-12 |
| 5 | 2016-01-01 | 2021-12-31 | 2015-07-01 | 2021-07-01 |
| 6 | 2014-01-01 | 2020-03-31 | 2013-07-01 | 2019-09-30 |
| 6I* | 2020-04-01 | 2021-12-31 | 2019-10-01 | 2021-07-01 |
| 6W* | 2020-04-01 | 2021-12-31 | 2019-10-01 | 2021-07-01 |

* This party split from party 6

### S2 Genetic and paternity analysis

Table S3: **Characteristics of the 24 microsatellite loci used to estimate paternity** (calculated using the genotypes of all males, females and offspring included in the analysis; $n = 256$)

| Loci | | Alleles | | Heterozygosity | | | NAFE | | Null alleles |
|---|---|---|---|---|---|---|---|---|---|
| Locus ID | Locus No | Allele range | No Alleles | He | Ho | HWE* | Brookfield | Chakraborty | Null alleles Presence |
| D6s264 | Locus1 | 94 – 98 | 3 | 0.521 | 0.506 | 0.891 | 0.010 | 0.015 | no |
| D7s503 | Locus2 | 152 – 166 | 6 | 0.756 | 0.784 | 0.213 | −0.016 | −0.018 | no |
| D12s375 | Locus3 | 165 – 181 | 5 | 0.742 | 0.753 | 0.793 | −0.006 | −0.007 | no |
| D3s1766 | Locus4 | 195 – 203 | 3 | 0.283 | 0.301 | 0.462 | −0.014 | −0.032 | no |
| D14s306 | Locus5 | 165 – 177 | 4 | 0.540 | 0.556 | 0.942 | −0.010 | −0.014 | no |
| D1s533 | Locus6 | 187 – 203 | 5 | 0.617 | 0.668 | 0.201 | −0.031 | −0.039 | no |
| D2s1329 | Locus7 | 212 – 228 | 5 | 0.628 | 0.595 | 0.794 | 0.021 | 0.027 | no |
| D2s1326 | Locus8 | 251 – 263 | 4 | 0.421 | 0.413 | 0.953 | 0.006 | 0.010 | no |
| D10s611 | Locus9 | 133 – 141 | 3 | 0.566 | 0.591 | 0.157 | −0.016 | −0.021 | no |
| D8s1106 | Locus10 | 144 – 160 | 5 | 0.467 | 0.494 | 0.951 | −0.018 | −0.028 | no |
| D17s791 | Locus11 | 166 – 172 | 4 | 0.571 | 0.602 | 0.443 | −0.020 | −0.026 | no |
| D6s501 | Locus12 | 172 – 192 | 6 | 0.691 | 0.749 | 0.075 | −0.034 | −0.040 | no |
| D17s1290 | Locus13 | 195 – 207 | 4 | 0.474 | 0.498 | 0.759 | −0.016 | −0.024 | no |
| D6s311 | Locus14 | 228 – 230 | 2 | 0.362 | 0.351 | 0.632 | 0.008 | 0.015 | no |
| D5s1457 | Locus15 | 124 – 136 | 4 | 0.374 | 0.398 | 0.785 | −0.017 | −0.031 | no |
| D8s505 | Locus16 | 147 – 151 | 2 | 0.285 | 0.282 | 0.810 | 0.002 | 0.005 | no |
| D10s1432 | Locus17 | 158 – 170 | 7 | 0.745 | 0.707 | 0.305 | 0.022 | 0.026 | no |
| D5s820 | Locus18 | 179 – 199 | 6 | 0.770 | 0.730 | 0.051 | 0.023 | 0.027 | no |
| D3s1768 | Locus19 | 169 – 213 | 6 | 0.554 | 0.541 | 0.083 | 0.009 | 0.012 | no |
| D7s2204 | Locus20 | 228 – 248 | 6 | 0.724 | 0.695 | 0.030 | 0.017 | 0.021 | no |
| D1s207 | Locus21 | 133 – 135 | 2 | 0.495 | 0.510 | 0.720 | −0.010 | −0.015 | no |
| D4s243 | Locus22 | 147 – 167 | 6 | 0.638 | 0.649 | 0.005 | −0.006 | −0.008 | no |
| D1s548 | Locus23 | 192 – 208 | 5 | 0.738 | 0.718 | 0.294 | 0.011 | 0.014 | no |
| D21s1142 | Locus24 | 226 – 246 | 6 | 0.697 | 0.722 | 0.464 | −0.015 | −0.018 | no |
| Mean | | | 4.5 | 0.569 | 0.575 | 0.492 | −0.004 | −0.006 | |
| SD | | | 1.5 | 0.151 | 0.149 | 0.339 | 0.017 | 0.022 | |
| Min | | | 2.0 | 0.283 | 0.282 | 0.005 | −0.034 | −0.040 | |
| Max | | | 7.0 | 0.770 | 0.784 | 0.953 | 0.023 | 0.027 | |

He= expected heterozygosity; Ho= observed heterozygosity; HWE= Hardy-Weinberg equilibrium (* note that $p$-value was corrected for multiple testing with the Bonferroni adjustment, $\alpha = (0.05/24) = 0.00208$)); NAFE= null alleles frequencies estimators calculated based on Brookfield (1996) and Chakraborty et al. (1992) and presence of null alleles.

Table S4: **Results of the paternity analysis for offspring born during the study period (2014-2021) in the study parties.** Samples and genotypes were available for a total of 71 offspring, while for the other 39 offspring no genetic information were available (see the column 'Offspring sampling'). Results from the paternity analysis conducted with Cervus 3.0 (version 3.0.7; Kalinowski et al., 2007). Nmis indicates number of mismatches; Trio LOD indicates the scores of the logarithm of the likelihood ratio; trio Delta is defined as the difference in LOD scores between the most likely and the second most likely candidate father. The confidence level of the Cervus paternity assignments was set to 95% ('strict' criterion) and all statistical confidence on paternity assignment were higher than 95%. An asterisk in the most likely father column indicates fathers that were not the unit holder at the time of conception. In particular, one asterisk indicate a father belonging to the same party and two asterisks a father belonging to a different party of the same gang. For two offspring the unit holder at time of conception was unknown (scored as NA) due to data gaps and the covid pandemic.

| Party | Offspring | Date of birth | Mother | Mother genetically confirmed | Offspring sampling | Most likely father | Unit holder at time of conception | $N_{\text{mis}}$ | Trio LOD | Trio Delta | Conf-level |
|---|---|---|---|---|---|---|---|---|---|---|---|
| 6 | O1 | 2014-01 | F10 | Y | Yes | M17* | M20 | 0 | 1.4e+15 | 1.3e+15 | >95% |
| 9 | O2 | 2014-03 | F27 | Y | Yes | M11 | M11 | 0 | 1.1e+15 | 1.5e+14 | >95% |
| 6 | O3 | 2014-04 | F11 | Y | Yes | M35 | M35 | 0 | 1.6e+15 | 1.6e+15 | >95% |
| 6 | O4 | 2014-06 | F26 | Y | Yes | M20 | M20 | 0 | 1.0e+15 | 1.0e+15 | >95% |
| 6 | O5 | 2014-07 | F19 | Y | Yes | M35 | M35 | 0 | 1.0e+15 | 2.7e+13 | >95% |
| 9 | O6 | 2014-07 | F41 | Y | Yes | M42 | M42 | 0 | 1.2e+15 | 1.2e+15 | >95% |
| 9 | O7 | 2014-08 | F16 | Y | Yes | M49 | M49 | 0 | 1.2e+15 | 6.7e+14 | >95% |
| 9 | O8 | 2014-09 | F47 | Y | Yes | M05 | M05 | 0 | 1.2e+15 | 7.0e+11 | >95% |
| 6 | O9 | 2014-09 | F44 | | No | | M51 | | NA | NA | |
| 9 | O10 | 2014-10 | F14 | Y | Yes | M11 | M11 | 1 | 6.7e+14 | 6.7e+14 | >95% |
| 9 | O11 | 2014-10 | F29 | | No | | M62 | | NA | NA | |
| 6 | O12 | 2015-01 | F45 | Y | Yes | M51 | M51 | 0 | 1.2e+15 | 9.8e+14 | >95% |
| 9 | O13 | 2015-02 | F28 | Y | Yes | M49 | M49 | 0 | 9.0e+14 | 9.0e+14 | >95% |
| 9 | O14 | 2015-03 | F39 | Y | Yes | M31 | M31 | 0 | 1.4e+15 | 1.4e+15 | >95% |
| 9 | O15 | 2015-04 | F12 | Y | Yes | M31 | M31 | 0 | 1.4e+15 | 1.2e+15 | >95% |
| 9 | O16 | 2015-05 | F23 | | No | | M62 | | NA | NA | |
| 9 | O17 | 2015-05 | F40 | Y | Yes | M63** | M49 | 0 | 8.5e+14 | 8.5e+14 | >95% |
| 6 | O18 | 2015-05 | F44 | Y | Yes | M51 | M51 | 0 | 1.2e+15 | 4.1e+14 | >95% |
| 6 | O19 | 2015-05 | F46 | Y | Yes | M35 | M35 | 0 | 1.2e+15 | 1.1e+15 | >95% |
| 6 | O20 | 2015-06 | F3 | Y | Yes | M28 | M28 | 0 | 1.2e+15 | 1.2e+15 | >95% |
| 9 | O21 | 2015-07 | F27 | | No | | M05 | | NA | NA | |
| 6 | O22 | 2015-07 | F7 | | No | | M56 | | NA | NA | |
| 9 | O23 | 2015-08 | F31 | Y | Yes | M62 | M62 | 0 | 1.0e+15 | 8.9e+14 | >95% |
| 9 | O24 | 2015-12 | F43 | Y | Yes | M05 | M05 | 0 | 6.7e+14 | 6.7e+14 | >95% |
| 5 | O25 | 2016-03 | F2 | Y | Yes | M38 | M38 | 0 | 1.4e+15 | 1.4e+15 | >95% |
| 5 | O26 | 2016-03 | F32 | Y | Yes | M43 | M43 | 0 | 1.3e+15 | 1.3e+15 | >95% |
| 9 | O27 | 2016-04 | F29 | Y | Yes | M62 | M62 | 0 | 1.6e+15 | 1.6e+15 | >95% |
| 9 | O28 | 2016-05 | F15 | Y | Yes | M05 | M05 | 0 | 9.7e+14 | 9.1e+14 | >95% |
| 6 | O29 | 2016-05 | F26 | | No | | M20 | | NA | NA | |
| 6 | O30 | 2016-05 | F10 | Y | Yes | M15* | M20 | 0 | 1.3e+15 | 1.3e+15 | >95% |
| 9 | O31 | 2016-05 | F8 | Y | Yes | M62 | M62 | 0 | 1.2e+15 | 9.6e+14 | >95% |
| 9 | O32 | 2016-05 | F25 | Y | Yes | M31 | M31 | 0 | 1.6e+15 | 1.6e+15 | >95% |
| 6 | O33 | 2016-05 | F19 | | No | | M35 | | NA | NA | |
| 6 | O34 | 2016-06 | F7 | Y | Yes | M21 | M21 | 0 | 1.1e+15 | 1.4e+14 | >95% |
| 9 | O35 | 2016-07 | F16 | Y | Yes | M49 | M49 | 0 | 1.4e+15 | 1.4e+15 | >95% |
| 6 | O36 | 2016-07 | F11 | Y | Yes | M35 | M35 | 0 | 8.9e+14 | 8.9e+14 | >95% |
| 9 | O37 | 2016-07 | F33 | Y | Yes | M62 | M62 | 0 | 1.1e+15 | 1.1e+15 | >95% |
| 9 | O38 | 2016-08 | F41 | | No | | M42 | | NA | NA | |
| 6 | O39 | 2016-08 | F45 | Y | Yes | M17 | M17 | 0 | 1.4e+15 | 5.3e+14 | >95% |
| 9 | O40 | 2016-09 | F14 | | No | | M42 | | NA | NA | |
| 9 | O41 | 2016-09 | F47 | Y | Yes | M05 | M05 | 0 | 1.3e+15 | 1.3e+15 | >95% |
| 6 | O42 | 2017-02 | F46 | | No | | M59 | | NA | NA | |
| 5 | O43 | 2017-02 | F48 | Y | Yes | M43 | M43 | 0 | 9.4e+14 | 9.1e+14 | >95% |
| 5 | O44 | 2017-02 | F20 | Y | Yes | M38 | M38 | 0 | 1.2e+15 | 1.2e+15 | >95% |
| 9 | O45 | 2017-03 | F40 | Y | Yes | M49 | M49 | 0 | 1.1e+15 | 1.1e+15 | >95% |
| 6 | O46 | 2017-03 | F44 | Y | Yes | M28 | M28 | 0 | 1.1e+15 | 1.1e+15 | >95% |
| 9 | O47 | 2017-03 | F12 | | No | | M31 | | NA | NA | |
| 6 | O48 | 2017-03 | F3 | Y | Yes | M28 | M28 | 0 | 1.3e+15 | 1.3e+15 | >95% |

Table S4 – continued from previous page

| Party | Off-spring | Date of birth | Mother | Mother genetically confirmed | Off-spring sam-pling | Most likely fa-ther | Unit holder at time of conception | $N_{\text{mis}}$ | Trio LOD | Trio Delta | Conf-level |
|---|---|---|---|---|---|---|---|---|---|---|---|
| 5 | O49 | 2017-03 | F37 | Y | Yes | M10* | M27 | 0 | 1.3e+15 | 1.0e+15 | >95% |
| 9 | O50 | 2017-05 | F28 | Y | Yes | M49 | M49 | 0 | 1.1e+15 | 1.1e+15 | >95% |
| 9 | O51 | 2017-05 | F23 | | No | | M62 | | NA | NA | |
| 5 | O52 | 2017-05 | F17 | | No | | M38 | | NA | NA | |
| 5 | O53 | 2017-06 | F13 | | No | | M08 | | NA | NA | |
| 9 | O54 | 2017-06 | F21 | Y | Yes | M31 | M31 | 0 | 1.2e+15 | 1.2e+15 | >95% |
| 6 | O55 | 2017-07 | F22 | | No | | M21 | | NA | NA | |
| 9 | O56 | 2017-09 | F31 | | No | | M62 | | NA | NA | |
| 5 | O57 | 2017-09 | F32 | | No | | M43 | | NA | NA | |
| 6 | O58 | 2017-11 | F46 | Y | Yes | M55 | M55 | 0 | 1.4e+15 | 1.4e+15 | >95% |
| 9 | O59 | 2018-01 | F43 | | No | | M05 | | NA | NA | |
| 6 | O60 | 2018-03 | F10 | | No | | M21 | | NA | NA | |
| 5 | O61 | 2018-04 | F13 | Y | Yes | M08 | M08 | 0 | 1.3e+15 | 1.3e+15 | >95% |
| 9 | O62 | 2018-04 | F25 | | No | | M42 | | NA | NA | |
| 5 | O63 | 2018-05 | F4 | Y | Yes | M08 | NA | 0 | 1.6e+15 | 1.6e+15 | >95% |
| 9 | O64 | 2018-05 | F16 | | No | | M49 | | NA | NA | |
| 6 | O65 | 2018-06 | F35 | Y | Yes | M15 | M15 | 0 | 9.1e+14 | 9.1e+14 | >95% |
| 6 | O66 | 2018-07 | F11 | Y | Yes | M21 | M21 | 0 | 8.6e+14 | 4.4e+14 | >95% |
| 9 | O67 | 2018-08 | F29 | Y | Yes | M62 | M62 | 0 | 1.3e+15 | 1.3e+15 | >95% |
| 5 | O68 | 2018-08 | F32 | | No | | M43 | | NA | NA | |
| 5 | O69 | 2018-10 | F1 | Y | Yes | M08 | M08 | 0 | 1.2e+15 | 1.2e+15 | >95% |
| 9 | O70 | 2018-10 | F12 | | No | | M31 | | NA | NA | |
| 9 | O71 | 2018-11 | F47 | Y | Yes | M05 | M05 | 0 | 1.1e+15 | 1.1e+15 | >95% |
| 6 | O72 | 2018-11 | F44 | Y | Yes | M28 | M28 | 0 | 1.2e+15 | 8.6e+14 | >95% |
| 6 | O73 | 2018-11 | F22 | | No | | M21 | | NA | NA | |
| 5 | O74 | 2019-01 | F2 | Y | Yes | M38 | M38 | 0 | 1.3e+15 | 1.3e+15 | >95% |
| 6 | O75 | 2019-02 | F42 | | No | | M04 | | NA | NA | |
| 5 | O76 | 2019-04 | F20 | | No | | M38 | | NA | NA | |
| 5 | O77 | 2019-04 | F9 | | No | | M55 | | NA | NA | |
| 6 | O78 | 2019-06 | F30 | Y | Yes | M61* | M28 | 0 | 1.4e+15 | 1.4e+15 | >95% |
| 6 | O79 | 2019-06 | F10 | Y | Yes | M21 | M21 | 1 | 1.1e+15 | 5.4e+14 | >95% |
| 5 | O80 | 2019-07 | F5 | Y | Yes | M10 | M10 | 0 | 1.7e+15 | 1.7e+15 | >95% |
| 6 | O81 | 2019-07 | F38 | Y | Yes | M21 | M21 | 0 | 1.1e+15 | 6.8e+14 | >95% |
| 5 | O82 | 2019-07 | F48 | Y | Yes | M43 | M43 | 0 | 1.1e+15 | 7.5e+14 | >95% |
| 5 | O83 | 2019-07 | F37 | | No | | M16 | | NA | NA | |
| 5 | O84 | 2019-10 | F46 | Y | Yes | M55 | M55 | 0 | 9.5e+14 | 9.5e+14 | >95% |
| 9 | O85 | 2020-02 | F8 | | No | | M53 | | NA | NA | |
| 6 | O86 | 2020-02 | F42 | | No | | M04 | | NA | NA | |
| 9 | O87 | 2020-02 | F28 | | No | | M02 | | NA | NA | |
| 6 | O88 | 2020-02 | F35 | | No | | M04 | | NA | NA | |
| 9 | O89 | 2020-02 | F40 | | No | | M40 | | NA | NA | |
| 9 | O90 | 2020-02 | F21 | | No | | M31 | | NA | NA | |
| 5 | O91 | 2020-02 | F13 | Y | Yes | M08 | M08 | 0 | 8.4e+14 | 7.3e+14 | >95% |
| 5 | O92 | 2020-03 | F36 | | No | | M55 | | NA | NA | |
| 5 | O93 | 2020-04 | F4 | Y | Yes | M08 | M08 | 0 | 8.1e+14 | 8.1e+14 | >95% |
| 5 | O94 | 2020-06 | F18 | Y | Yes | M55 | M55 | 0 | 1.4e+15 | 1.4e+15 | >95% |
| 5 | O95 | 2020-06 | F1 | Y | Yes | M08 | M08 | 0 | 1.8e+15 | 1.8e+15 | >95% |
| 5 | O96 | 2020-07 | F37 | Y | Yes | M08 | M08 | 0 | 1.3e+15 | 1.3e+15 | >95% |
| 6W | O97 | 2020-09 | F11 | Y | Yes | M21 | M21 | 0 | 1.1e+15 | 1.0e+15 | >95% |
| 6W | O98 | 2020-10 | F44 | Y | Yes | M28 | M28 | 0 | 1.3e+15 | 1.3e+15 | >95% |
| 6W | O99 | 2021-01 | F38 | Y | Yes | M21 | M21 | 0 | 1.3e+15 | 1.0e+15 | >95% |
| 6W | O100 | 2021-02 | F24 | Y | Yes | M28 | M28 | 0 | 9.0e+14 | 9.0e+14 | >95% |
| 5 | O101 | 2021-02 | F2 | Y | Yes | M38 | M38 | 0 | 1.1e+15 | 1.1e+15 | >95% |
| 6I | O102 | 2021-02 | F34 | Y | Yes | M04 | NA | 0 | 1.6e+15 | 1.3e+15 | >95% |
| 5 | O103 | 2021-03 | F46 | Y | Yes | M55 | M55 | 0 | 9.6e+14 | 9.6e+14 | >95% |
| 6W | O104 | 2021-03 | F30 | Y | Yes | M28 | M28 | 1 | 1.1e+15 | 1.1e+15 | >95% |
| 6W | O105 | 2021-07 | F10 | Y | Yes | M21 | M21 | 0 | 1.3e+15 | 1.2e+15 | >95% |
| 5 | O106 | 2021-07 | F32 | | No | | M55 | | NA | NA | |
| 6I | O107 | 2021-08 | F35 | | No | | M04 | | NA | NA | |
| 5 | O108 | 2021-09 | F13 | Y | Yes | M08 | M08 | 0 | 9.8e+14 | 7.8e+14 | >95% |
| 5 | O109 | 2021-10 | F1 | | No | | M08 | | NA | NA | |
| 6W | O110 | 2021-10 | F6 | | No | | M21 | | NA | NA | |

### S3 Reproductive skew

We calculated reproductive skew using the Multinomial Skew Index ($M$ index, [1]).

When we fitted the model by [1] to our data, we ran into severe convergence issues. We therefore reparameterized the model and adapted the Stan code accordingly. In essence, we changed the parameter that reflected the individual probabilities in the multinomial model from a constrained vector (simplex) to an unconstrained vector that we fit as normally distributed. With this modification, the model ran without convergence issues on our data. In figure S1, we visualize posteriors of $M$ that come from model fits using both model parameterizations using the six data sets on human populations from [1]. This figure shows that our modification seems to be by and large equivalent to the original implementation.

We also subjected our modified model to a simple set of posterior predictive checks. For this we used the 50 posterior draws of the estimated probability vector to predict actual numbers of offspring sired by each male. Figure S2 shows the results of this exercise. Within each male, the bubbles indicate the expectation for how many offspring were sired when predicted from the model. When comparing the model predictions to the observed data, we can see that model makes on average adequate predictions.

In addition to calculating $M$ from the observed data, we also calculated $M$ for two hypothetical boundary scenarios [2]. To arrive at the first hypothetical $M$ when a male monopolizes all reproduction $M_{\text{monopoly}}$, we created a data set where one male sired all offspring, while the other males sired 0 offspring. Accordingly, $M_{\text{equal}}$ comes from a synthetic data set where each male obtained exactly one offspring (i.e. equal share). Both scenarios were adapted for each party (table 1 in main text).

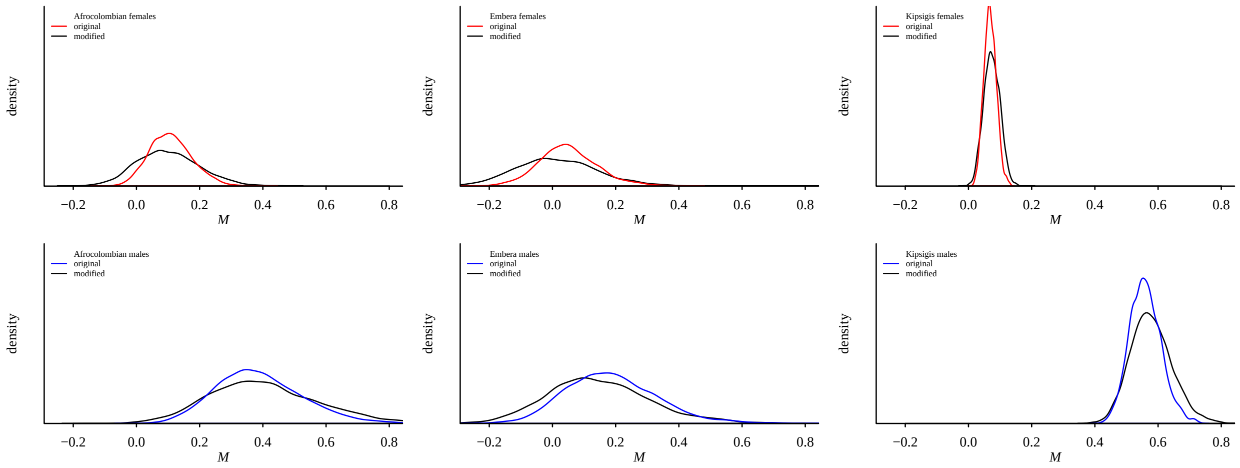


Figure S1: Comparison between the original and modified algorithms to arrive at posterior distributions of the M index (Ross et al 2020). Fits are from six data sets (see figure 2 in Ross et al 2020). In general, there is good agreement between the two algorithms.

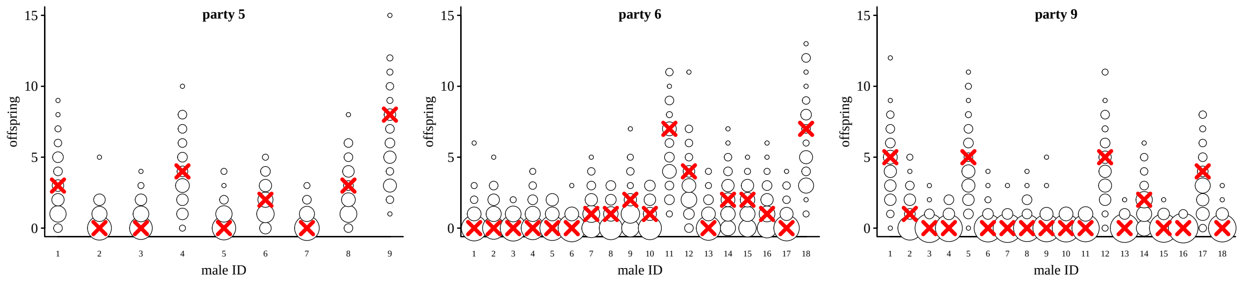


Figure S2: Posterior predictive check for the reproductive skew models. Here we compare predictions from 50 random posterior draws (circles) with respect to the number of offspring a male sired. The observed values are the red crosses. Circle size is proportional to the predicted number of sired offspring.

### S4 Modelling potential reproductive success

This section contains a more detailed model description and some additional model checks and extended results. We fitted our models in Stan 2.33.1 [3] with the cmdstanr R interface [v. 0.6.1.9000, 4] in R 4.6.1 [5].

Below we show a detailed specification of our model. For ease of display, we split the description into two parts. The first part describes the modeling of Elo-ratings. The second part describes the actual modeling of the number of females a male had in his unit.

#### S4-1 Elo-ratings

We started with declaring a $j+1$ by $s$ matrix, where $j$ is the number of observed dyadic dominance interactions and $s$ is the number of subjects (males). The first row in this matrix corresponded to the ratings of all individuals before any interactions were observed, i.e., it represents the vector of start ratings.

$$M = \begin{pmatrix} \text{elo}_{\text{start,subject[1]}} & \cdots & \text{elo}_{\text{start,subject[s]}} \\ \text{elo}_{1,\text{subject[1]}} & \cdots & \text{elo}_{1,\text{subject[s]}} \\ \cdots & \cdots & \cdots \\ \text{elo}_{j,\text{subject[1]}} & \cdots & \text{elo}_{j,\text{subject[s]}} \end{pmatrix} \tag{S1}$$

The remaining rows of $M$ were filled iteratively, going through each interaction in turn. We also declared a vector $p_{\text{win}}$, in which we stored the winning probabilities for each interaction.

$$p_{\text{win}} = \begin{pmatrix} p_{\text{win},1} & \cdots & p_{\text{win},j} \end{pmatrix} \tag{S2}$$

In the first step of the iterative Elo-rating algorithm, ratings were centered among those individuals who were present in the group at the date of the observed interaction $j$.

Second, we calculated the expected winning probability $p_{\text{win},j}$ of the subject that won the interaction. This step required only the ratings of the winning and the losing subjects *prior* to the interaction ($\text{M}_{j-1,\text{subject[winner}[j]]}$ and $\text{M}_{j-1,\text{subject[loser}[j]]}$). In the case of the first interaction in the sequence, these ratings were the start ratings.

$$p_{\text{win},j} = \frac{1}{1 + exp(\text{M}_{j-1,\text{loser}[j]} - \text{M}_{j-1,\text{winner}[j]})} \tag{S3}$$

With the obtained winning probability, we updated the ratings of the winning and losing individuals in $M_j$, and keep the ratings of individuals that were not involved in interaction $j$, like so:

$$M_{j,s} = \begin{cases} M_{j-1,s} + (1 - p_{\text{win},j}) * k & \text{if } s \text{ won the interaction} \\ M_{j-1,s} - (1 - p_{\text{win},j}) * k & \text{if } s \text{ lost the interaction} \\ M_{j-1,s} & \text{otherwise} \end{cases} \tag{S4}$$

We then repeated the steps in equations S3 and S4 over all observed interactions, and filled $M$ in this way. Most importantly, we obtained the vector $p_{\text{win}}$ (with the expected winning probabilities), which we then used as the probability vector to model the actual observed outcomes as Bernoulli trials (see [6]). Up to this point the algorithm is deterministic (for a given combination of start ratings and $k$).

To estimate the two required quantities (start ratings and $k$), we then fitted the following model using $p_{\text{win}}$ (which in turn depended on the start ratings and $k$ (equations S3 and S4)). The response in this model was simply a vector of length $j$, filled with 1s. This vector encoded that the winner of an interaction did indeed win the interaction [6]. We set priors for the start ratings and $k$ following [6].

$$
\begin{aligned}
\text{win}_j &\sim \text{Bernoulli}(p_{\text{win},j}) \quad j = \text{interactions } 1...714 \\
\text{elo}_{start} &\sim \text{Normal}(0, 1) \\
k &\sim \text{Exponential}(2)
\end{aligned}
\tag{S5}
$$

Once we estimated the vector of start ratings and $k$, we obtained the rating of subject $s$ at a given date. This rating represented the predictor value in the subsequent count model. To arrive at such a rating, we ran the deterministic Elo-rating algorithm to obtain $M$, using the estimated start ratings and $k$. Since we were interested in ratings at the end of calendar years, we looked up the last row in $M$ where the corresponding interaction date belonged to the year of interest. Note that these ratings were not point estimates, but full posterior distributions (see figure 2 in main text). In other words, each rating we used in the count model was a posterior distribution, which incorporated uncertainty in the estimation of a subject's rating at a given time.

In the description of the actual count model, we refer to this step of obtaining the rating for a given observation via the function $F()$.

Note that we treated the five groups/parties separately, i.e., we estimated five $k$ values and five start rating vectors. For simplicity, we illustrated our model design up to this point with one group only.

#### S4-2 Count model

$$
y_i \sim \text{Poisson}(\lambda_i) \qquad i = \text{observations } 1...190 \tag{S6}
$$

We used a Poisson likelihood and log-link function, and as predictors we used a male's Elo-rating, age and the interaction between Elo-rating and age.

$$
log(\lambda_i) = \mathcal{A}_i + \mathcal{B}_i * \text{elo}_i + \mathcal{C}_i * \text{age}_i + \mathcal{D}_i * \text{elo}_i * \text{age}_i + \epsilon_i \tag{S7}
$$

As described above, we estimated start ratings and $k$ to arrive at male ratings that corresponded to a given observation.

$$
\text{elo}_i = F(\text{elo}_{start}, k, \text{interactions}, \text{year}[i], \text{subject}[i]) \tag{S8}
$$

To account for overdispersion, we used an observation-level varying intercept ($\epsilon$) [7].

$$
\begin{aligned}
\epsilon &\sim \text{Normal}(0, \sigma_e) \\
\sigma_e &\sim \text{Exponential}(5)
\end{aligned}
\tag{S9}
$$

We fitted varying intercepts and correlated varying slopes for all terms. Specifically, we incorporated varying effects for Elo-rating and age in all grouping variables (subject, group/party, year). For the interaction between age and Elo-rating, we incorporated varying slopes only on the level of group/party and year, but not for males because less than half of the males who were present in the data (20 out of 48) changed age category over the course of the study and only 4 males had at least two data points in each age category.

$$
\begin{aligned}
\mathcal{A}_i &= \alpha + \alpha_{\text{subject}[i]} + \alpha_{\text{group}[i]} + \alpha_{\text{year}[i]} && \text{intercept} \\
\mathcal{B}_i &= \beta + \beta_{\text{subject}[i]} + \beta_{\text{group}[i]} + \beta_{\text{year}[i]} && \text{slope for Elo-rating} \\
\mathcal{C}_i &= \gamma + \gamma_{\text{subject}[i]} + \gamma_{\text{group}[i]} + \gamma_{\text{year}[i]} && \text{slope for age} \\
\mathcal{D}_i &= \delta + \delta_{\text{group}[i]} + \delta_{\text{year}[i]} && \text{slope for interaction}
\end{aligned} \tag{S10}
$$

$$
\begin{aligned}
\begin{bmatrix} \alpha_{\text{subject}} \\ \beta_{\text{subject}} \\ \gamma_{\text{subject}} \end{bmatrix} &\sim \text{MVNormal}\left( \begin{bmatrix} 0 \\ 0 \\ 0 \end{bmatrix}, S_{\text{s}} \right) && \text{subject} = 1...48 \\
S_{\text{s}} &= \begin{pmatrix} \sigma_{sa} & 0 & 0 \\ 0 & \sigma_{sb} & 0 \\ 0 & 0 & \sigma_{sc} \end{pmatrix} R_s \begin{pmatrix} \sigma_{sa} & 0 & 0 \\ 0 & \sigma_{sb} & 0 \\ 0 & 0 & \sigma_{sc} \end{pmatrix} \\
\begin{bmatrix} \alpha_{\text{group}} \\ \beta_{\text{group}} \\ \gamma_{\text{group}} \\ \delta_{\text{group}} \end{bmatrix} &\sim \text{MVNormal}\left( \begin{bmatrix} 0 \\ 0 \\ 0 \\ 0 \end{bmatrix}, S_{\text{g}} \right) && \text{group} = 1...5 \\
S_{\text{g}} &= \begin{pmatrix} \sigma_{ga} & 0 & 0 & 0 \\ 0 & \sigma_{gb} & 0 & 0 \\ 0 & 0 & \sigma_{gc} & 0 \\ 0 & 0 & 0 & \sigma_{gd} \end{pmatrix} R_g \begin{pmatrix} \sigma_{ga} & 0 & 0 & 0 \\ 0 & \sigma_{gb} & 0 & 0 \\ 0 & 0 & \sigma_{gc} & 0 \\ 0 & 0 & 0 & \sigma_{gd} \end{pmatrix} \\
\begin{bmatrix} \alpha_{\text{year}} \\ \beta_{\text{year}} \\ \gamma_{\text{year}} \\ \delta_{\text{year}} \end{bmatrix} &\sim \text{MVNormal}\left( \begin{bmatrix} 0 \\ 0 \\ 0 \\ 0 \end{bmatrix}, S_{\text{y}} \right) && \text{year} = 1...9 \\
S_{\text{y}} &= \begin{pmatrix} \sigma_{ya} & 0 & 0 & 0 \\ 0 & \sigma_{yb} & 0 & 0 \\ 0 & 0 & \sigma_{yc} & 0 \\ 0 & 0 & 0 & \sigma_{yd} \end{pmatrix} R_y \begin{pmatrix} \sigma_{ya} & 0 & 0 & 0 \\ 0 & \sigma_{yb} & 0 & 0 \\ 0 & 0 & \sigma_{yc} & 0 \\ 0 & 0 & 0 & \sigma_{yd} \end{pmatrix}
\end{aligned} \tag{S11}
$$

For all parameters, we used fairly vague priors (see figure S3):

$$
\begin{aligned}
R_s &\sim \text{LKJ}(2) \\
\sigma_{sa}, \sigma_{sb}, \sigma_{sc} &\sim \text{Exponential}(5) \\
R_g &\sim \text{LKJ}(2) \\
\sigma_{ga}, \sigma_{gb}, \sigma_{gc}, \sigma_{gd} &\sim \text{Exponential}(5) \\
R_y &\sim \text{LKJ}(2) \\
\sigma_{ya}, \sigma_{yb}, \sigma_{yc}, \sigma_{yd} &\sim \text{Exponential}(5) \\
\alpha, \beta, \gamma, \delta &\sim \text{Normal}(0, 1)
\end{aligned} \tag{S12}
$$

For eight males (representing eleven male-years) that were included in the count model, we had no observed dominance interactions. Those males entered the data set for the count model with ratings based simply on the specified prior for the start ratings (equation S5).

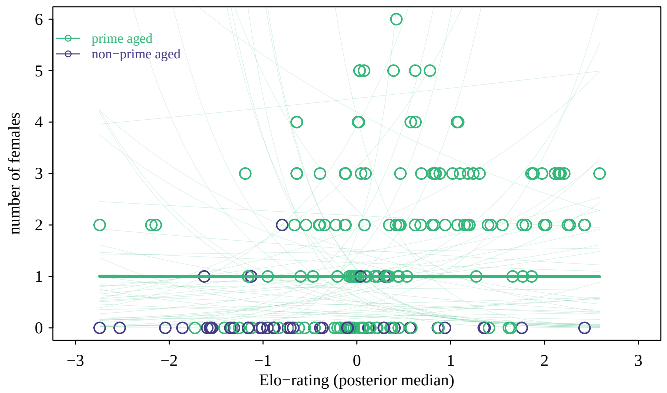


Figure S3: Prior predictive simulation. This plot illustrates the potential range of model estimates for the primary parameter of interest (Elo), given our priors. The lines show 100 individual predictions. On average the priors encode a flat relationship between Elo-rating and number of females (thick line). However, the priors allow negative as well as positive relationships. Note also that our priors lead to a substantial proportion of predictions that are unrealistic, i.e., predicting 10s or 100s of females and therefore could be set tighter (more informative) in future iterations of the model. Note also that for this simulation the parameters related to Elo-rating ($k$ and start rating vectors) were estimated from the data.

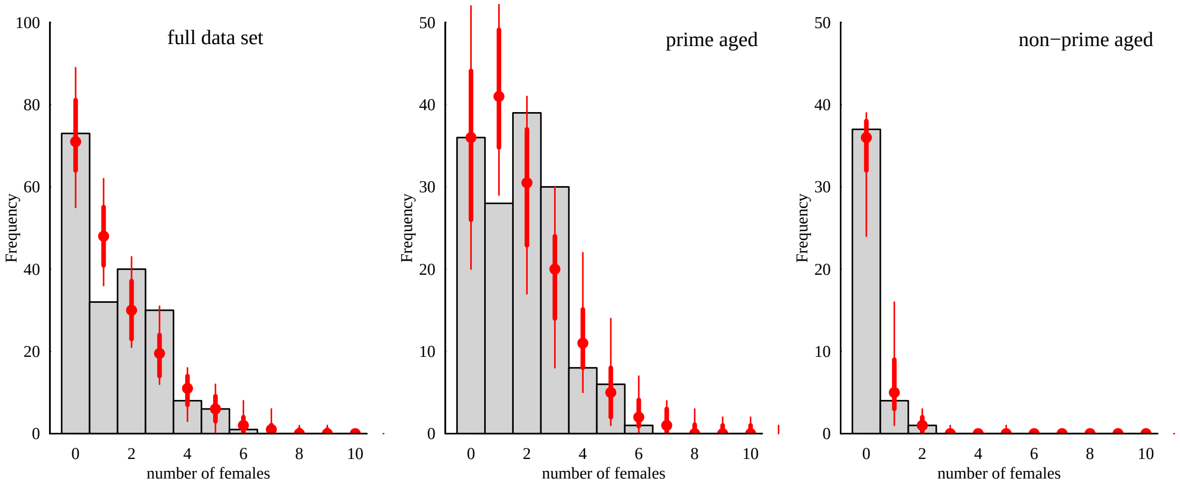


Figure S4: Posterior predictive checks. The histogram shows the observed distribution of number of females. The circles represent the median of 50 randomly selected random draws. The thick lines represent 80% quantiles and the thin lines represent the range across the 50 draws. The figure shows that our model overestimates the occurrences of males having one female compared to what we observed, and underestimates the occurrences of males having 3 females. The two remaining plots show the same approach, but separated by the two age categories.

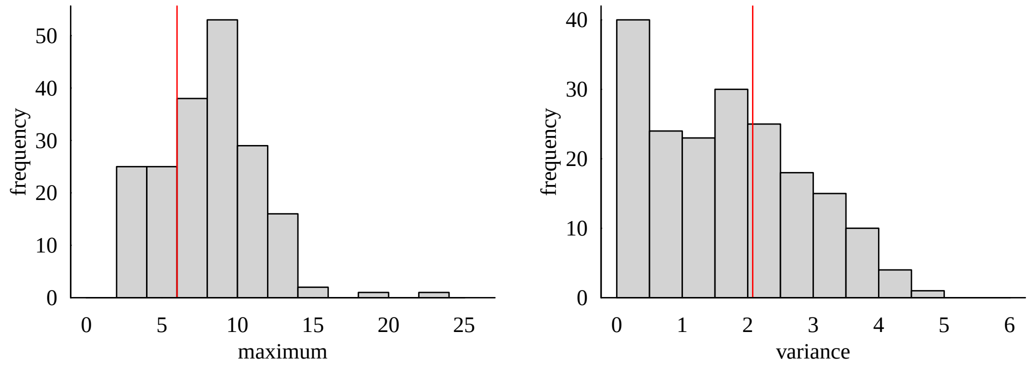


Figure S5: Additional posterior predictive checks. The histograms show the posterior distributions of two summary statistics applied to all posterior samples: maximum value of females and variance in the number of females. The red vertical lines show the corresponding values from the observed data. This plot shows that our model performs well with respect to predicting these two aspects of our observed data.

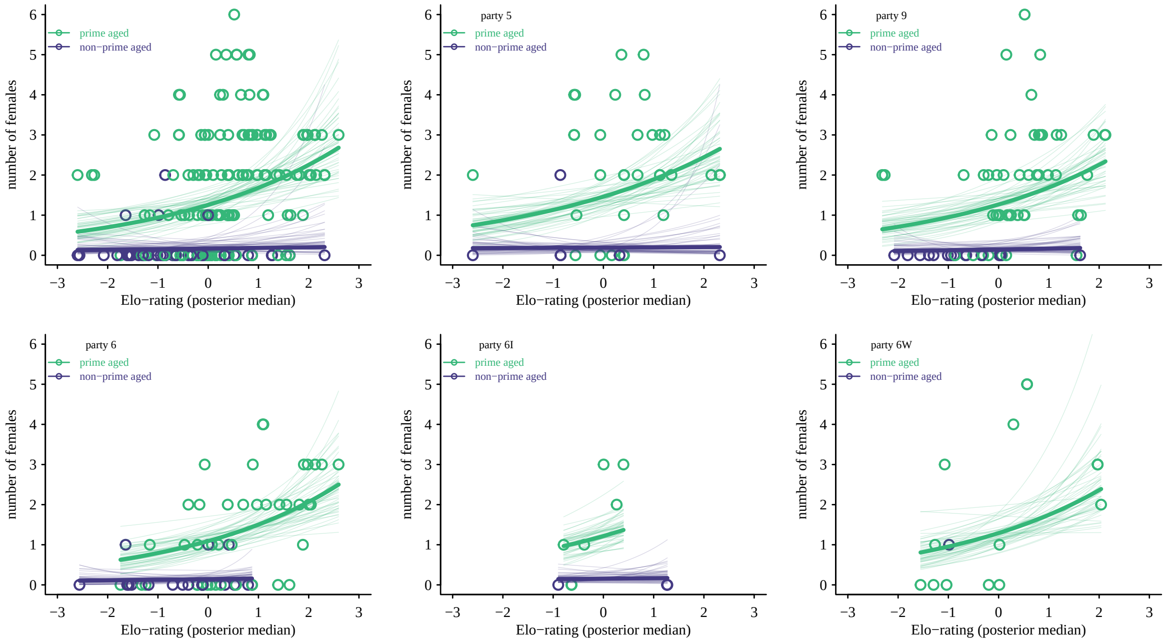


Figure S6: Same as figure 3 in main text, but split by party. There was suprisingly little variation across the parties in the estimated relationship between Elo-rating and number of females.

Table S5: Extended results for the reproductive success model.

| | mean | median | 5.5% | 94.5% | $\hat{R}$ | $\text{ESS}_{\text{bulk}}$ | $\text{ESS}_{\text{tail}}$ |
|---|---|---|---|---|---|---|---|
| intercept ($\alpha$) | 0.23 | 0.23 | -0.04 | 0.48 | 1.00 | 2074 | 2524 |
| Elo-rating ($\beta$) | 0.29 | 0.29 | 0.10 | 0.49 | 1.00 | 1988 | 2519 |
| age ($\gamma$) | -2.02 | -2.00 | -2.73 | -1.33 | 1.00 | 4089 | 2738 |
| interaction Elo-rating : age ($\delta$) | -0.21 | -0.21 | -0.78 | 0.38 | 1.00 | 3219 | 2748 |
| **varying intercepts (SD)** | | | | | | | |
| male ID, $\sigma_{sa}$ | 0.46 | 0.45 | 0.28 | 0.65 | 1.00 | 1392 | 1459 |
| group/party, $\sigma_{ga}$ | 0.18 | 0.16 | 0.02 | 0.42 | 1.00 | 1406 | 1803 |
| year, $\sigma_{ya}$ | 0.08 | 0.06 | 0.01 | 0.20 | 1.00 | 2473 | 2300 |
| **varying slopes (Elo-rating) (SD)** | | | | | | | |
| male ID, $\sigma_{sb}$ | 0.10 | 0.08 | 0.01 | 0.26 | 1.00 | 1311 | 1761 |
| group/party, $\sigma_{gb}$ | 0.10 | 0.07 | 0.01 | 0.27 | 1.00 | 2412 | 2331 |
| year, $\sigma_{yb}$ | 0.07 | 0.05 | 0.01 | 0.18 | 1.00 | 2017 | 1737 |
| **varying slopes (age) (SD)** | | | | | | | |
| male ID, $\sigma_{sc}$ | 0.21 | 0.15 | 0.01 | 0.59 | 1.00 | 2670 | 1731 |
| group/party, $\sigma_{gc}$ | 0.21 | 0.15 | 0.01 | 0.62 | 1.00 | 3336 | 2495 |
| year, $\sigma_{yc}$ | 0.19 | 0.14 | 0.01 | 0.54 | 1.00 | 4134 | 2143 |
| **varying slopes (interaction) (SD)** | | | | | | | |
| group/party, $\sigma_{gd}$ | 0.17 | 0.12 | 0.01 | 0.48 | 1.00 | 3214 | 1569 |
| year, $\sigma_{yd}$ | 0.20 | 0.14 | 0.01 | 0.56 | 1.00 | 2759 | 2099 |
